# No reproductive fitness benefits of dear enemy behaviour in a territorial songbird

**DOI:** 10.1101/2021.04.28.441816

**Authors:** Michael S. Reichert, Jodie M.S. Crane, Gabrielle L. Davidson, Eileen Dillane, Ipek G. Kulahci, James O’Neill, Kees van Oers, Ciara Sexton, John L. Quinn

**Affiliations:** Department of Integrative Biology, Oklahoma State University, USA; School of Biological Earth and Environmental Sciences, University College Cork, Ireland; Kākāpō Recovery Programme, Department of Conservation, Invercargill, New Zealand; Department of Psychology, University of Cambridge, Cambridge, UK; Behavioural Ecology Group, Wageningen University, Wageningen, The Netherlands; Department of Animal Ecology, Netherlands Institute of Ecology (NIOO-KNAW), Wageningen, The Netherlands

**Keywords:** habituation, individual recognition, playback, great tit, territorial behaviour, cognition

## Abstract

Territorial animals often respond less aggressively to neighbours than strangers. This ‘dear enemy’ effect is hypothesized to be adaptive by reducing unnecessary aggressive interactions with non-threatening individuals. A key prediction of this hypothesis, that individual fitness will be affected by variation in the speed and the extent to which individuals reduce their aggression towards neighbours relative to strangers, has never been tested. We used a series of song playbacks to measure the change in response of male great tits to a simulated establishment of a neighbour on an adjacent territory during early stages of breeding, as an assay of individuals’ tendencies to form dear enemy relationships. Males reduced their approach to the speaker and sang fewer songs on later playback repetitions. However, only some males exhibited dear enemy behaviour by responding more strongly to a subsequent stranger playback, and when the playback procedure was repeated on a subset of males, there was some indication for consistent differences among individuals in the expression of dear enemy behaviour. We monitored nests and analysed offspring paternity to determine male reproductive success. Individuals that exhibited dear enemy behaviour towards the simulated neighbour did not suffer any costs associated with loss of paternity, but there was also no evidence of reproductive benefits, and no net effect on reproductive fitness. The general ability to discriminate between neighbours and strangers is likely adaptive, but benefits are probably difficult to detect because of the indirect link between individual variation in dear enemy behaviour and reproductive fitness, and because of the complex range of mechanisms affecting relations with territorial neighbours.

## INTRODUCTION

Territoriality is a widespread behaviour that provides important benefits to territory holders such as reducing competition for food or breeding resources, and guarding of mates from extra-pair matings (Stamps 1994; Adams 2001). However, there are many costs to holding territories that arise primarily from the vigilance and aggressive behaviours required to defend and maintain territorial boundaries against intruders (Ydenberg 1984; Mares et al. 2012; Wischhoff et al. 2018). One way to lower these costs is to reduce territorial defence behaviours towards non-threatening individuals and instead focus defence efforts particularly on those individuals that pose a threat of usurping the territory or the resources within (Getty 1987; Temeles 1994). Thus, many species exhibit the ‘dear enemy effect’, which is defined as individuals showing less aggressive behaviour towards territorial neighbours than towards strangers (Ydenberg et al. 1988; Temeles 1994; Stoddard 1996; Tumulty 2018). If neighbours are not a threat because they hold their own territory, then dear enemy behaviour should benefit individual fitness through reduced costs of unnecessary aggression, facilitating greater investment in foraging and reproduction (Getty 1987; Temeles 1994), as well as defence against truly threatening individuals (Leiser and Itzkowitz 1999). This hypothesis predicts that intraspecific variation in the speed and extent to which individuals reduce their aggression towards non-threatening neighbours will affect individual fitness, particularly enhancing reproductive success when the territory is used for breeding. However, this key prediction has never been tested.

Selection acts on individual variation, and thus to understand the evolution of the dear enemy effect it is important to determine whether individuals differ consistently in dear enemy behaviour (as a proxy for heritability), and whether these differences have fitness consequences. Repeatable individual variation in dear enemy effect expression is expected because of variation in individuals’ cognitive capabilities to recognize familiar neighbours (Reichert and Quinn 2017), because of personality traits such as aggressiveness (Akçay et al. 2014), and due to covariance with other repeatable behavioural traits (Verbeek et al. 1996). Some studies have provided indirect support for fitness benefits of dear enemy behaviour, for instance by demonstrating that individuals holding territories with long-term neighbours have higher reproductive success than those with new neighbours, or that neighbours tend to engage in cooperative behaviours together (Beletsky and Orians 1989; Grabowska-Zhang et al. 2012a, b; Siracusa et al. 2021). However, in these studies it is unknown whether and to what extent individuals discriminated between neighbours and strangers in their aggressive interactions, and therefore whether there was variation in dear enemy behaviour. Thus, while it is clear that the ability and tendency to discriminate neighbours from strangers varies and likely has functional significance, there is still a limited understanding of the selection pressures that may be acting on dear enemy behaviour, especially at the within-species level.

The dear enemy effect is facilitated by cognitive mechanisms that enable individuals to learn some characteristic of their neighbours, allowing them to discriminate between neighbours and strangers (Wiley 2013). These mechanisms – of which more than one may be involved in any given species – range from habituation to a neighbour’s signal characteristics and/or their spatial location (Petrinovich and Peeke 1973; Brooks and Falls 1975; Yasukawa 1981; Bee and Gerhardt 2001; Dong and Clayton 2009), associative learning of neighbour characteristics (Richards 1979), to ‘true individual recognition’ (Johnston and Jernigan 1994; Gheusi et al. 1997; Saeki et al. 2018), and act over time scales ranging from short-term decreases in aggression within a day to persistent recognition of individuals across years (Godard 1991; Tumulty and Sheehan 2020). Thus, variation among individuals in the expression of the dear enemy effect (i.e. the extent and speed with which aggression is reduced towards neighbours relative to strangers) may arise because of variation in the cognitive abilities associated with neighbour-stranger discrimination. If this cognitive variation affects reproductive success—for instance if those individuals on breeding territories that quickly learn to recognize their neighbours avoid unnecessary agonistic encounters and can invest more in their offspring—then individuals with superior cognitive ability may have higher fitness and be favoured by selection. Non-cognitive factors also affect the expression of the dear enemy effect. For instance, individuals may differ in aggressiveness or other personality traits (Hyman et al. 2004; Amy et al. 2010; Jacobs et al. 2014), and context-dependent factors such as territory size, breeding status, and the density of neighbouring territory-holders also likely affect responsiveness to neighbours (Werba et al. 2021). Regardless of the source, this individual variation in dear enemy expression has potentially significant, but unexplored, consequences for fitness. Furthermore, few studies have examined the trajectory over which the dear enemy effect develops upon establishment of a new neighbour (e.g., Bee and Gerhardt 2001), or whether this differs among individuals.

Like many songbirds, great tits (*Parus major*) defend territories around their nest during the breeding season (Gosler 1993). Male great tits do so by singing and approaching intruding individuals to engage in visual displays and occasionally, physical fighting (Blurton Jones 1968; Krebs 1977). Great tits nest in natural cavities or artificial nest boxes, and the primary threat from intruders is the potential for them to usurp the limited resource of a high-quality nesting location (Krebs 1971, 1976, 1982), although intruders also present other threats including potentially mating with the territory-holder’s social mate (Hill et al. 2011) or foraging on the territory (Hinde 1956). In great tits, approximately 25-50% of broods contain extrapair young (Lubjuhn et al. 1999; Brommer et al. 2010; Patrick et al. 2012). The dear enemy effect has been demonstrated previously in great tits: in playback experiments, individuals showed a reduced response to songs of their territorial neighbours compared to songs of strangers (Krebs 1971; Falls et al. 1982; McGregor and Avery 1986), and the effect was stronger when the neighbour’s song was played from its own territory (McGregor and Avery 1986). Territory defence begins very quickly after settlement, often within hours (Krebs 1971). The speed with which dear enemy relationships are formed is unknown, but given that strong territory defence occurs during a limited breeding season, and that great tits habituate rapidly to song (Krebs 1976), a reduced response to neighbours compared to strangers is likely to occur within a few days.

Recognition of neighbours in great tits is not based on a simple discrimination between the categories of familiar and unfamiliar individuals (Wiley 2013); instead they can learn to discriminate among the songs of different specific individuals (McGregor and Avery 1986; Weary and Krebs 1992). There is also among-individual variation in aggressive responses to playback resulting from personality differences (Amy et al. 2010), and personality and reproductive investment were related to the speed with which individuals habituated to a series of song playbacks (Rivera-Gutierrez et al. 2017); this habituation to a familiar stimulus is likely an important mechanism in the establishment of dear enemy relations (Peeke 1984). Great tits trade off the time invested in foraging and territory defence (Ydenberg 1984; Ydenberg and Krebs 1987); thus, individuals that rapidly reduce their aggression towards their neighbours may benefit directly by increasing their foraging intake, and this in turn could enhance their ability to provision for their offspring (Martin 1987). However, vigilance must be maintained against potential territorial usurpers, because the longer a usurper is on the territory, the more effort is required to expel it (Krebs 1982).

We performed a series of acoustic playback experiments in which we monitored territorial males’ responses to the simulated arrival of a new male on a neighbouring territory. Our overall objective was to examine whether individuals habituated by reducing their response after repeated exposures to the songs of a simulated new neighbour while still maintaining a heightened response to a stranger (i.e., displayed dear enemy behaviour), whether they differed consistently in this regard, and whether there were any beneficial effects of the tendency to form dear enemy relationships on fitness and reproduction in general. We note that we did not expect these fitness effects to arise because of the individual’s actual behaviours towards the simulated neighbour in our playback trials, but rather we assumed that the individual’s behaviour in the playback trials was indicative of how it engaged with others in natural territorial interactions, with variation in the tendency to reduce aggression towards a familiar stimulus (as an experimental proxy for the tendency to form dear enemy relationships) leading to effects on reproductive success.

First, we determined if males’ responses to playbacks of a simulated neighbour that were broadcast multiple times across three days declined over time, consistent with habituation to the neighbour playback stimulus. Second, we tested whether the response to a subsequent ‘stranger’ playback was stronger than that to the final neighbour playback, consistent with the reduced aggressive response to a familiar neighbour compared to a stranger that is the hallmark of the dear enemy effect. Third, we estimated the repeatability of several dear enemy behaviour measures to determine if individuals differed intrinsically in these behaviours. We also explored the extent to which the expression of these dear enemy behaviour measures was context dependent, for example due to phenology, which is related to overall levels of nest defence (Hyman 2005; Jin et al. 2021), and the repetition rate of songs during the playback, which affects the rate of habituation towards repeated stimuli (Thompson et al. 1973). Finally, we tested whether there was any evidence for selection or any reproductive benefits of dear enemy behaviour in a number of ways: A) controlling for paternity, we tested the prediction of positive selection on the tendency to perform dear enemy behaviour, that is, that individuals that rapidly reduced their response to neighbours should fare better for a variety of life history traits that are good indicators of reproductive fitness and recruitment to the breeding population in the great tit (Tinbergen and Boerlijst 1990), in particular clutch size, the number of offspring fledged, and average offspring mass; B) We repeated these analyses including all nestlings (including extra-pair) to test for evidence of any benefits of the tendency to perform dear enemy behaviour at the nest, irrespective of paternity; C) We examined if there was any relationship between observed levels of extra-pair paternity at the social nest and the tendency to perform dear enemy behaviour. We discuss the implications of our experimental results and analyses of reproductive success for our understanding of the dear enemy effect.

## METHODS

Playbacks were performed during the spring breeding season (April-May in 2017 and 2018) in eight small forestry plots in County Cork, Ireland (Table S1, Fig. S1). Each site contained an array of nestboxes, which are preferentially used by great tits for breeding (East and Perrins 1988), and in which most individuals had been ringed as part of a long-term study (for details see O’Shea et al. 2018). We identified potential playback subjects by listening for males singing near nestboxes and examining the progress of nest building in the box. Males chosen for the experiment were then subject to a series of song playbacks as described below.

**Table 1.** Tests of change in response to neighbour playback across trials. Output is from a generalized linear mixed model for response (binary, yes or no), a cumulative link mixed model for closest approach (distances were placed into ordinal categories), and a zero-inflated glmm for number of songs (Poisson; the coefficients are for the conditional model output). The reference category for stimulus rate is the low rate (estimate = 0). Trial number was entered as a numerical variable, with values from one (first trial with neighbour stimulus) to nine (last trial with neighbour stimulus). *N* = 51 individuals

| Variable | Factor | Estimate (SE) | z | P |
| --- | --- | --- | --- | --- |
| Response | Intercept | 0.53 (0.27) |  |  |
|  | Trial number | -0.10 (0.04) | -2.52 | 0.01 |
|  | Rate (high) | 0.04 (0.28) | 0.13 | 0.89 |
| Closest approach | Trial number | 0.11 (0.04) | 3.10 | 0.002 |
|  | Rate (high) | -0.01 (0.30) | -0.03 | 0.97 |
| Number of songs | Intercept | 2.91 (0.09) |  |  |
|  | Trial number | -0.04 (0.01) | -5.88 | <0.001 |
|  | Rate (high) | 0.23 (0.12) | 1.89 | 0.058 |

### Playback stimuli

Male great tits sing on territories containing their nesting site. They typically have a repertoire of three to four distinctive song types (Gompertz 1961; McGregor and Krebs 1982a) but usually repeat the same one for several minutes before switching to a different type (Krebs 1976). The playbacks were designed to mimic this repetition of a single song type. Although simulating a larger song repertoire through playbacks may have captured additional aspects of the dear enemy phenomenon, this was not necessary to address our primary aims and would have reduced our power to detect dear enemy behaviour because: 1. Males habituate to the presentation of both a single song type and multiple song types, but habituation is slower to playback of multiple song types (Krebs 1976), and 2. Males will respond to the presentation of a single song type in a manner consistent with the dear enemy effect: responding more strongly to a single song type of a stranger than to a single song type of an established neighbour (Krebs 1971), 3. Great tits are capable of discriminating between individuals even based on songs of those individuals they have never heard before (Weary and Krebs 1992). Thus, our comparison of birds’ responses to a simulated neighbour (recorded song from one individual, played back several times) and a simulated stranger (recorded song from a different individual, played back once) is an appropriate assay of dear enemy behaviour (i.e., the difference in response to a familiar stimulus from a familiar location, the neighbour, and an unfamiliar stimulus from an unfamiliar location, the stranger) and is unlikely to have been perceived by the bird as two different song types from the same individual.

The playback stimuli consisted of recordings of natural male songs made using Wildlife Acoustics SM4 audio recorders (24 kHz sampling rate) placed at nestboxes in May 2016. We scanned audio files for exemplars of male songs with a high signal to noise ratio and containing no other bird songs in the background. We inserted each chosen song into a new audio file in Audacity software, bandpass filtered the song between 1.5 and 11.5 kHz, and manipulated the song to contain 6 phrases (the basic repeating unit of the song (McGregor and Krebs 1982b)) by copying or deleting phrases as needed. The song exemplar was then copied so that it was played back at a rate of either five or ten songs per minute for five minutes, which we refer to as the “low” and “high” stimulus rate treatments, respectively. These two stimulus rates were used to test the prediction that the likelihood of habituation to the neighbour songs depended on the stimulus repetition rate (Thompson and Spencer 1966). A total of 29 song exemplars were chosen, and each exemplar was recorded at a different nestbox; songs came from seven of the eight study sites. The playback stimulus selected for the subject male and the song rate treatment were chosen randomly, with the restriction that the stimulus song was not recorded from the same site as the subject (all sites were at least 2.25 km apart from each other; Table S1). This ensured that subjects would not have been familiar with the playback song already. Stimuli were broadcast as .wav files from an EasyAcc X02s speaker mounted on a tripod at approximately 1 m height, at a sound-pressure level of 90 dB (A) measured at 1 m using an Extech 407730 sound level meter. Great tit males respond readily to playback of song exemplars by exhibiting territorial behaviour (Rivera-Gutierrez et al. 2015; Snijders et al. 2017; Jin et al. 2021).

### Experimental design

On each of three consecutive days we exposed males to three playbacks, performed at 1-hour intervals, of one of the playback stimuli (Fig. 1). Each subject was therefore exposed to a total of nine playbacks of the same stimulus (henceforth the “neighbour” stimulus), which simulated a newly arrived male on a neighbouring territory. Territory settlement generally takes place very rapidly in this species (Krebs 1971), and responses to repeated playback typically decrease over a few days (Krebs et al. 1981; Rivera-Gutierrez et al. 2017), so this interval allowed us to study the initial development of dear enemy behaviour. The playbacks were always performed from the same location, and at a distance of 25 m from the nestbox. This location was chosen because territoriality is likely strongest near the nestbox (Giraldeau and Ydenberg 1987), 50 m is a typical nearest-neighbour distance (Krebs 1971), and neighbour-stranger discrimination is typically strongest near the territorial border (Falls and Brooks 1975; Stoddard et al. 1991). There is an inevitable trade-off between the advantages of our approach of broadcasting playbacks from a standard amplitude and distance from the nestbox, and those of estimating males’ actual territorial boundaries and placing the speaker at each male’s estimated territorial edge. Because we used a fixed playback distance, our playbacks may have been more salient to some individuals than others, depending on whether the standard 25 m playback distance was or was not within their territory. However, although great tits differ in their response to playbacks well outside of their territory [10 m beyond the boundary] and those very close to the centre of their territory [20-30 m within the boundary] (Peake et al. 2001), our setup was unlikely to have corresponded to this situation, and to our knowledge no study has examined whether great tits are capable of discriminating between songs coming from just inside versus just outside territorial boundaries. In contrast, it is well established that song intensity affects the aggressive response of songbirds, including great tits (Brumm and Ritschard 2011; Ritschard et al. 2012; Luther et al. 2016). Furthermore, song amplitude is likely to play a role in the speed of dear enemy learning (Bee 2001). We therefore used a fixed playback distance so that song amplitude could be standardized. The specific location of the playback was chosen in a randomized direction, with the constraint that playbacks were only performed from areas the experimenter could access, and that did not overlap with the territory of another male great tit.

**Fig. 1.**
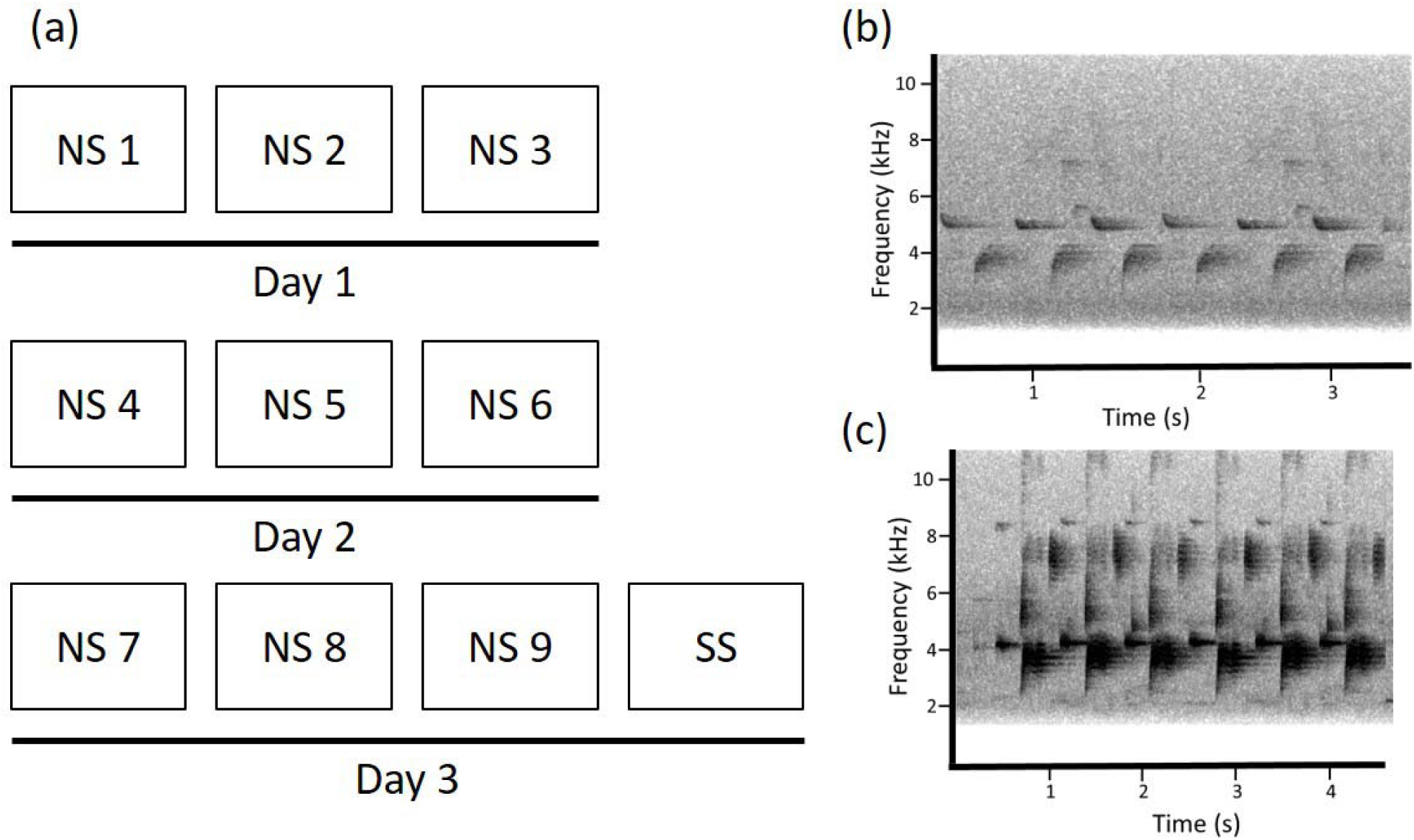
(a) Timeline of the playback experimental design. We broadcast the neighbour stimulus (‘NS’) for five minutes per trial (number in each box corresponds to the trial number), for a total of three trials per day with approximately one hour between trials. This procedure was repeated for three consecutive days. On the third day, immediately following the final neighbour playback (NS 9), we broadcast a different song from a different location (‘SS’: stranger stimulus, a test of stimulus specificity, in other words that males that had reduced their response to the neighbour retain the capability of an aggressive response towards strangers). In 2017, the entire procedure was repeated with different stimuli after a period of ten days to test the repeatability of dear enemy behaviour. (b) Exemplar playback stimulus. The spectrogram shows a single great tit song that was used to build one of the playback stimuli. (c) a second exemplar stimulus, taken from a different bird, illustrating the acoustic variation in great tit songs. An example playback experiment could use the song in (b) for the neighbour stimulus and that in (c) for the stranger stimulus. Note that the spectrograms do not illustrate the equalization of playback amplitude that was used for the trials

On the third day, immediately after the response was recorded to the ninth exposure of the subject to the neighbour stimulus, we broadcast a different song stimulus from a different location (still 25 m away from the nestbox, and at a 90-degree angle or greater, with respect to the nest box, from the location of the neighbour playbacks). The aim was to compare the response to this new stimulus, simulating a new stranger individual (henceforth, the “stranger” stimulus), to the original playback stimulus to which we hypothesized the subject would have developed some familiarity towards. This is an essential step for demonstrating neighbour-stranger discrimination in territorial systems, in which individuals reduce their response to familiar neighbours, but maintain a heightened response to an unfamiliar stranger (Bee et al. 2016). In other words, the stranger playback was used to demonstrate that the decline in aggressive response observed across the trials is specific to the neighbour’s song. We performed the stranger playback immediately after recording the response to the last neighbour playback for logistical reasons and because doing so is a more conservative test of neighbour-stranger discrimination than if we had delayed the stranger playback by some interval. If an individual was not responsive to its neighbour but then immediately responded to the stranger playback, this is a strong test of the ability to discriminate between the two, whereas any delay between the playbacks increases the chance that any generalized habituation of the aggressive response would decline, potentially leading to misinterpretation of the data. Although it is possible that birds could have mounted an increased aggressive response during the stranger playback session not because they were responding to what they perceived as a stranger but rather because it was a novel situation in that the playback occurred a few minutes, rather than an hour, after the previous playback, we consider this interpretation unlikely because for this to be the case either birds would have had to learn the number of daily playbacks or the time interval between them, or would have had to interpret the playback as a different song type of the neighbour, which is unlikely because great tits are capable of learning individual voice characteristics and then discriminating among them even based on songs they had never heard before (Weary and Krebs 1992) and therefore should not have mistaken our stranger stimulus for another song of the neighbour.

We justify our design of using both a different location and song for the stranger stimulus, because the aim of our experiment was not to identify the specific criteria used by individuals to discriminate neighbours and strangers, i.e., whether discrimination was based on these individuals’ song characteristics or location, but rather to determine the consequences of individual differences in tendency to discriminate. Therefore, we used a playback design that increased the opportunities for discrimination between our stimuli by varying both location and song, which is also the most likely scenario for natural neighbour-stranger discrimination because the neighbour would not normally move to a different territory and the stranger would not normally sing from the exact spot that the neighbour was just singing in. We acknowledge that our design does not separate the effects of location and song type by including controls for each. This would not have been feasible given the longitudinal design and limited breeding window in this wild system. Furthermore, although we cannot rule out that subjects interpreted a song type recorded from a different bird and broadcast from a different location as in fact still coming from the original simulated neighbour, as discussed above, great tits discriminate among individual voice characteristics (Weary and Krebs 1992). Therefore, we interpret their responses in the final playback as responses to a simulated stranger, but note that, even under dear enemy relationships, individuals may also respond aggressively to neighbours when they display from a new location (Brindley 1991; Husak and Fox 2003; Lovell and Lein 2005; Dalton et al. 2020).

In 2017 only, we repeated this three-day procedure with 20 individuals ten days after their final trial from the first round of playbacks. The aim of this repetition was to investigate whether the expression of dear enemy behaviour was repeatable, which would indicate intrinsic differences among individuals (Bell et al. 2009). Therefore, we performed playbacks as above, but with a different song stimulus from a different location, simulating a different new neighbour establishing a territory. A different stranger stimulus was also used in these repeated trials, and the male was given the alternative rate treatment to that which it had been exposed during the first round of playbacks.

Playbacks were performed between 0800h and 1540h; each male was tested at approximately the same time on each of the three days of the experiment. Playbacks occurred during the early stages of breeding for most birds – between late nest building and egg laying. Due to logistical constraints of running experiments across multiple field sites with respect to the timing of nest checks, seven individuals were tested after the last egg was laid at the nest, including three individuals for which incubation was already started at the nest (the latest start date of the first repetition of the playback trials was 4 days into incubation). To control for the timing of the playback with respect to the breeding cycle (Petrinovich et al. 1976; Mace 1987; Jin et al. 2021), we included the date of the first playback trial relative to the first egg date as a variable in analyses (see below).

### Playback procedure

During each playback a single observer was positioned in cover near the playback speaker to monitor the male’s behaviour, noting the distance between the bird and the speaker, and making audio recordings of any songs using Marantz PMD 660 or PMD 661 audio recorders with Sennheiser ME67 directional microphones (16 bit, 44.1 kHz sampling rate). We did our best to avoid being observed by the birds during the playback; we consider it unlikely that our presence affected the results because these experiments took place in areas close to busy walking trails and birds were likely habituated to the presence of humans. All playbacks took place in locations where a male was known to occupy a territory because on the days prior to the playbacks it was observed singing near the nestbox and there was evidence of nest building activity. Males were usually visually or acoustically located prior to the start of the playback, and if not, most were sighted during the playback. This was not always possible, however, and we assume that these males initially unseen were within hearing distance of the stimulus during the playback. This was reasonable because males spend the great majority of their time on their territories and when they do leave their territories they only move short distances (Firth et al. 2018); at such distances our playback stimulus amplitude would remain well above the species’ masked hearing threshold (Langemann et al. 1998). Nevertheless, it is possible that we did not detect some response vocalizations from birds that were located far from the recorder, although signal to noise ratios of great tit song remain high at least 60 m from the source (Blumenrath and Dabelsteen 2004; Mockford et al. 2011). Joint territory defence by two individuals in response to the playback was not observed in this study, possibly due to the low density of individuals at our study sites. It was not possible to record data blind because our study involved focal animals in the field.

### Response variables

We noted the closest approach of the male to the playback speaker, in categories of 5 m (Nelson and Soha 2004). Subjects were not always located at the nest box at the start of the playback, so values greater than 25 m were possible. Individuals that did not respond at all were given a value of “None” for closest approach, which in the ordinal analyses described below was considered the greatest distance (whereas closest approaches from 0-5 m were considered the shortest distance, with successive distance categories ranked according to their distance). From the audio recordings we counted the number of songs produced by the male during the playback. We considered males to have responded to the playback if they sang at any point during the stimulus broadcast, or made any movement towards the playback speaker, and to have not responded if they did neither of those. The final dataset included 51 individuals; data from the first trial recording for one bird included in this total were lost because of a faulty microphone cable.

### Criterion for dear enemy behaviour

The typical method for demonstrating the dear enemy effect is to present individuals, in a single playback session, with the signals of an actual established neighbour, followed by a single playback session presenting the signals of an individual that the subject could not have interacted with previously (Brunton et al. 2008; Wei et al. 2011; Battiston et al. 2015). The dear enemy effect is inferred when individuals have a reduced response to the neighbour stimulus compared to the stranger stimulus. In our protocol we simulated the establishment of a new neighbour through a series of playbacks, which essentially served as training sessions to give the subject the opportunity to learn to recognize its neighbour. Therefore, to test whether or not the subject was indeed exhibiting dear enemy behaviour at the end of the necessarily fixed number of trials, we used the typical criterion for testing the dear enemy effect: comparison of the response to a final neighbour playback and a stranger playback. Our criterion (hereafter referred to as the ‘standard criterion’) that the subject was expressing dear enemy behaviour was that it did not respond in the final (ninth) playback of the simulated neighbour, but then did respond to the subsequent (tenth) playback of a simulated stranger.

Although our binary criterion should theoretically identify individuals that were expressing dear enemy behaviour towards the simulated neighbour, it has its limitations, so we explored additional measures of an individual’s change in response towards its neighbour over time. First, in some cases individuals responded to neither the final neighbour playback nor the stranger playback (N = 21). These results are difficult to interpret because they may indicate a general loss of motivation to defend the territory rather than a failure to discriminate neighbours and strangers. We therefore repeated all analyses, defining dear enemy behaviour as above, but only including those birds that responded to the stranger stimulus (referred to as the ‘standard criterion without non-responders’; N = 30 of 51 birds meeting this criterion; no individual failed to respond to the stranger playback after responding to the final neighbour playback). Second, for each individual, we extracted slope parameters from a logistic regression of the binary response variable on trial number (not including the stranger playback; referred to as the ‘response slope criterion’), and a linear regression of the number of songs on trial number (referred to as the ‘song slope criterion’). The aim was to obtain a more quantitative estimate of the change in response across trials that may reveal more variation than our binary criterion. If individuals have developed dear enemy behaviour towards the newly established neighbour, these values would be expected to be negative, indicating a decline in response across trials. However, a decline in response to a neighbour is not sufficient to demonstrate the dear enemy effect, because individuals must continue to respond to strangers. Therefore, we only included individuals that responded to the stranger playback for this second set of analyses (N = 30).

### Breeding data

Breeding data were obtained as part of standard monitoring protocols for the project (O’Shea et al. 2018). We recorded the date when the first egg was laid, the total clutch size, date of hatching and number of fledglings. The identity of the subject male sometimes could be determined by identification of unique colour rings if the male had already been captured prior to the experiment. Some males were also identified using RFID-equipped nestbox entrance doors, which could read the unique passive integrated transponder tag placed on the leg of previously captured males. The age (first year juvenile or adult) of previously captured males was determined from capture records. Males could also be identified when breeding adults were caught at the nest for ringing, ageing (as either first year juveniles or adults), and measurements, 10-12 days after the eggs hatched. Fifteen subjects were not identified because the nest was abandoned prior to trapping (N=1 before eggs laid, N=8 before eggs hatched, N=5 after eggs hatched), or the male could not be caught (N=1). However, our analyses do not rely on knowing the specific identity of the subject, and we can safely assume that no male was recorded in the study more than once based on the timing and distribution of boxes at which playbacks were carried out across the eight sites. Breeding densities are low at our sites and in cases where males could not be identified by colour-rings we nevertheless consider it highly unlikely that more than one individual responded to the playback on different trials. Chicks were weighed at day 15, and we determined the number of fledglings by inspecting the nest for any dead chicks after the breeding attempt was complete.

### Paternity analysis

Estimates of male reproductive fitness can be strongly influenced by extra-pair paternity (EPP) (Webster et al. 1995). Although rates of EPP are relatively low in great tits (van Oers et al. 2008; Patrick et al. 2012), they could have altered the relationship between dear enemy behaviour and reproductive fitness, particularly as paternity loss is one of the potential costs of territorial intrusions, and neighbours and strangers may differ in the threat they pose to paternity (Schlicht et al. 2015). EPP levels may also influence selection on males to cooperate via dear enemy effects (Eliassen and Jørgensen 2014). We therefore analyzed males’ reproductive success using metrics that excluded any offspring that were identified in a paternity analysis as being extra-pair offspring (note that we did not attempt to quantify males’ success at obtaining extra-pair matings at other nests because the small size and fragmented nature of our study sites prevented us from confidently assessing a male’s reproductive output away from his own nest). This is the appropriate method to analyse fitness, but we were also interested in the consequences of dear enemy behaviour on parental care behaviour. Therefore, we performed an additional set of analyses on the reproductive success variables in which both within-pair and extra-pair young were included, and present these in the supplement (results were not qualitatively affected by whether or not we excluded extra-pair young).

DNA was obtained from feathers taken from breeding pairs and offspring on their respective dates of capture and ringing. DNA extraction was performed using the protocol of the E.Z.N.A. Tissue DNA Kit (Omega Bio-Tek, Norcross, GA, USA), with the exception of the use of 80 µL elution buffer on a single elution step. Samples were genotyped at eight microsatellite loci, selected based on previously observed variability and utility, as well as potential for multiplexing in a single reaction (Pma69u (k=7 alleles) (Kawano 2003); PmaD22 (k=20), PmaCan1 (k=15), PmaGAn30 (k=5), PmaC25 (k=19), PmaTAGAn86 (k=19), PmaTGAn33 (k=17) and PmaTGAn45 (k=11) (Saladin et al. 2003)). Multiplex PCR was performed in a 3.5 µl total volume, including 1 µl of DNA extract, and 1.75 µl of 2x Top-Bio™ Plain Combi PP Mastermix, with a concentration of 0.03 µM for the Pma69U, PmaCan1, PmaGAn30 and PmaTAGAn86 primers and 0.06 µM for the PmaC25, PmaD22, PmaTGAn33 and PmaTGAn45 primers. The PCR was programmed with an initial denaturation at 95°C (15 min) followed by five cycles of 94°C (30 s), 55°C (90 s), 72°C (60 s), then 27 cycles at 94°C (30 s), 57°C (90 s), 72°C (60 s), followed by an elongation step at 60°C for 30 min. The PCR products were diluted in 14 µL nuclease free water; and run on an Applied Biosystems ABI3500xl DNA analyser using POP-7 polymer with GeneScan™ 600 LIZ™ Dye Size Standard v2.0 (ThermoFisher Scientific). We used GeneMarker version 2.7.0 software (SoftGenetics, Pennsylvania, USA) to determine allele sizes.

Paternity was assigned using CERVUS version 3.0.7 software (Kalinowski et al. 2007) using 10,000 cycles, 94 candidate fathers, a 0.02% error rate, two candidate parents and 93% of loci typed as simulation parameters. Only individuals that were successfully genotyped at five or more loci were included in paternity analyses. Individuals were determined to be within-pair offspring if all loci matched those of the social father and social mother combination, or if there was a mismatch at only one of the loci but the social father was identified as the most likely father using critical trio LOD scores returned by the program. Offspring that did not meet these criteria were categorized as extra-pair offspring. We assumed that offspring whose paternity was not determined (9 of 96 fledged offspring and 13 of 111 weighed offspring were of unknown paternity; two offspring fledged but were not weighed or analysed for paternity because they fledged on the day of weighing) were within-pair offspring. The combined exclusion probability for all eight microsatellites was >99.99%. Two of our loci significantly deviated from Hardy-Weinberg equilibrium when the genotypes of all individuals in the analysis were included (PmaC25: χ^2^ = 40.98, df = 15, p < 0.001, PmaD22: χ^2^ = 33.87, df = 15, p = 0.004). This was likely due to the family structure of the data.

### Data analysis

Except for the analyses of repeatability (see below), all analyses were performed only on the first repetition of the playback trials (i.e., we excluded data from the second repetition of the three-day procedure that was performed in 2017 only). We address the following questions in our analyses:

1. Does the response to the neighbour playback decrease over time? We tested whether males reduced their aggressiveness as they became more exposed to the songs of another male using three different variables, all of which have been demonstrated to be related to the aggressive response in great tits in previous studies (McGregor and Avery 1986; Doutrelant et al. 2000; Amy et al. 2010; Snijders et al. 2017): “response”, i.e. whether the male responded at all by singing or approaching (binary), closest approach (ordinal categorical), and number of songs during individual trials. In all cases the trial number (1-9) and stimulus rate treatment (low or high) were entered as fixed effects, and individual identity as a random effect. Response was modelled using a binomial generalized linear mixed model using the glmer function in the lme4 version 1.1-23 package (Bates et al. 2015) in R 4.0.2 software (R Development Core Team 2021). Approach category was modelled using a cumulative link mixed model to account for the ordinal nature of the dependent variable using the clmm function in the ordinal 2019.12-10 package (Christensen 2019). Number of songs was modelled as a poisson variable using the glmmTMB function of the glmmTMB 1.0.2.1 package (Brooks et al. 2017), which accounts for zero inflation in the dataset.
2. Is there a difference in the response to the stranger playback compared to the final (ninth) familiar neighbour playback? If there was a decrement in aggression in the analyses above, the next step to demonstrate dear enemy behaviour is to show that this decrement is specific to the neighbour stimulus. Therefore, individuals were predicted to respond more strongly to the stranger playback than to the final neighbour playback. We tested the same variables as above, using the same analyses but with the trial variable a factor with two levels: final neighbour or stranger playback trial.
3. Is the tendency for individuals to form a dear enemy relationship within the timeframe of our playback design repeatable? We estimated the repeatability of neighbour-stranger discrimination using the rptR package (Nakagawa and Schielzeth 2010) with the repetition (first or second) and stimulus rate treatment as fixed factors, individual as a random factor and whether the individual did or did not meet the standard criterion (i.e., responded to the stranger but not the neighbour stimulus) as a binary dependent variable. One individual was removed from this analysis because it did not respond during any trial in the second repetition. Repeatabilities of the response and song slope measures were analysed similarly, but with the dependent variable modelled as Gaussian. We examined the context-dependence of dear enemy behaviour using separate models for each of the four different methods of quantifying dear enemy behaviour (see *Criterion for dear enemy recognition*, above). We used generalized linear models for the standard criterion and standard criterion without non-responders, and linear models for the response slope and song slope. In all cases, we tested for effects of year (2017 or 2018), date of the first playback, date of the first playback minus the date the first egg was laid (to account for variation in dear enemy behaviour across the breeding cycle (Jin et al. 2021)), stimulus repetition rate (low or high), and the average number of songs the subject produced across the nine neighbour playbacks, as an estimate of its overall ‘aggressiveness’. We performed separate models for male age, with model structure as above but age (first year or adult) as the only factor, because age was only known for a subset of the birds.
4. Is there evidence for a difference in reproductive fitness between individuals that did or did not express dear enemy behaviour? We ran separate models for each of four different metrics of reproductive success, controlling for paternity, and each of the four methods of quantifying dear enemy behaviour (see *Criterion for dear enemy recognition*, above). For these analyses, we excluded data from three males from one of the sites (Dunderrow, see Table S1) in 2018 because of widespread nest predation. Although predation is certainly a component of reproductive success, in this site almost every nest was completely predated by stoats, *Mustela erminea*, and therefore we consider that there could be no relationship between the male phenotypic characteristics under study and the survival of offspring in this site with unusually high predation, rendering reproductive measures of these individuals meaningless in the context of our hypotheses. In all reproductive success models we included the metric of dear enemy behaviour, as defined above, as a fixed factor. Initial models also included the date the first egg was laid in the clutch, with the first of March as day 1, as well as the date of the first playback trial relative to the first egg date, but neither of these variables ever explained variation in reproductive success and so were excluded from the final models (similar results on the lack of effects of these variables on reproductive success were found by (O’Shea et al. 2018)). Four measures of reproductive success were analysed as follows: A) Clutch size was modelled as a Poisson variable in a generalized linear model. For one subject, no eggs were laid at the nestbox, and it was therefore excluded from all analyses of reproductive success; B) Number of within-pair fledglings was a Poisson variable, but there were a large number of zero values. We therefore modelled it with a glmmTMB model that accounts for zero inflation. In addition, because of the large number of zero values, we ran a model with the binary dependent variable of whether or not any offspring were fledged from the nest to compare males with successful versus unsuccessful nests; C) Average mass of within-pair offspring on day 15. Offspring biomass is an important determinant of fitness (Tinbergen and Boerlijst 1990), although this variable excludes the large number of subjects (N=18) whose nests failed entirely before day 15 (we had no evidence that these failures were caused by predation (e.g. broken eggs or predated remains), and instead were likely due to either parental abandonment or natural death of all offspring; note that the latter two causes are not readily distinguished), which of course is a severe fitness cost. We entered average offspring mass as the dependent variable in a linear model with an additional factor of brood size (including both within- and extra-pair offspring); D) The actual mass of each individual within-pair offspring. This analysis also excludes failed nests but may reveal important variation in parental investment among those nests that did survive. Individual mass was entered as a Gaussian variable in a linear mixed model, with brood size as an additional factor and nest as a random effect. Initial models with site included as a random effect could not be run because of singularity issues, because there were few samples from most sites (see Table S1). For the same reason, we did not perform a formal analysis of reproductive success by site, but there were no obvious qualitative differences in these variables between the four sites with the largest numbers of individuals tested (Table S1). We then repeated the above analyses including all nestlings (i.e., both within-pair and extra-pair) to determine if there was evidence for any benefit in the social nest of the tendency to perform dear enemy behaviour. We did not reanalyse clutch size because the analyses above also included all eggs, both within- and extra-pair, because we did not determine paternity for all eggs.
5. Finally, we tested for a relationship between dear enemy behaviour and extra-pair paternity using the proportion of extra-pair young at the nest as the dependent variable (binomial; general linear model), and, in separate models, each measure of dear enemy behaviour as a factor.

## RESULTS

### Change in response to the neighbour playback

The likelihood of a response (singing and/or moving towards the speaker) significantly decreased across the neighbour playback trials (Table 1; Fig. 2a). The pattern of responses shows an initial slight increase in the likelihood of responding over the first few trials, and then larger decreases especially in the final trials of the day. Closest approach was significantly greater on later trials such that individuals were more likely to make a close approach to the speaker on earlier trials, and approached but stayed further away or did not approach at all on later trials (Table 1; Fig. 2b). Likewise, the number of songs produced by males during the playback decreased across trials (Table 1; Fig. 2c). There was no effect of stimulus rate on response or closest approach, but there was a non-significant trend for an effect on the number of songs, with males giving more songs in response to stimuli presented at the higher rate (Table 1).

**Fig. 2.**
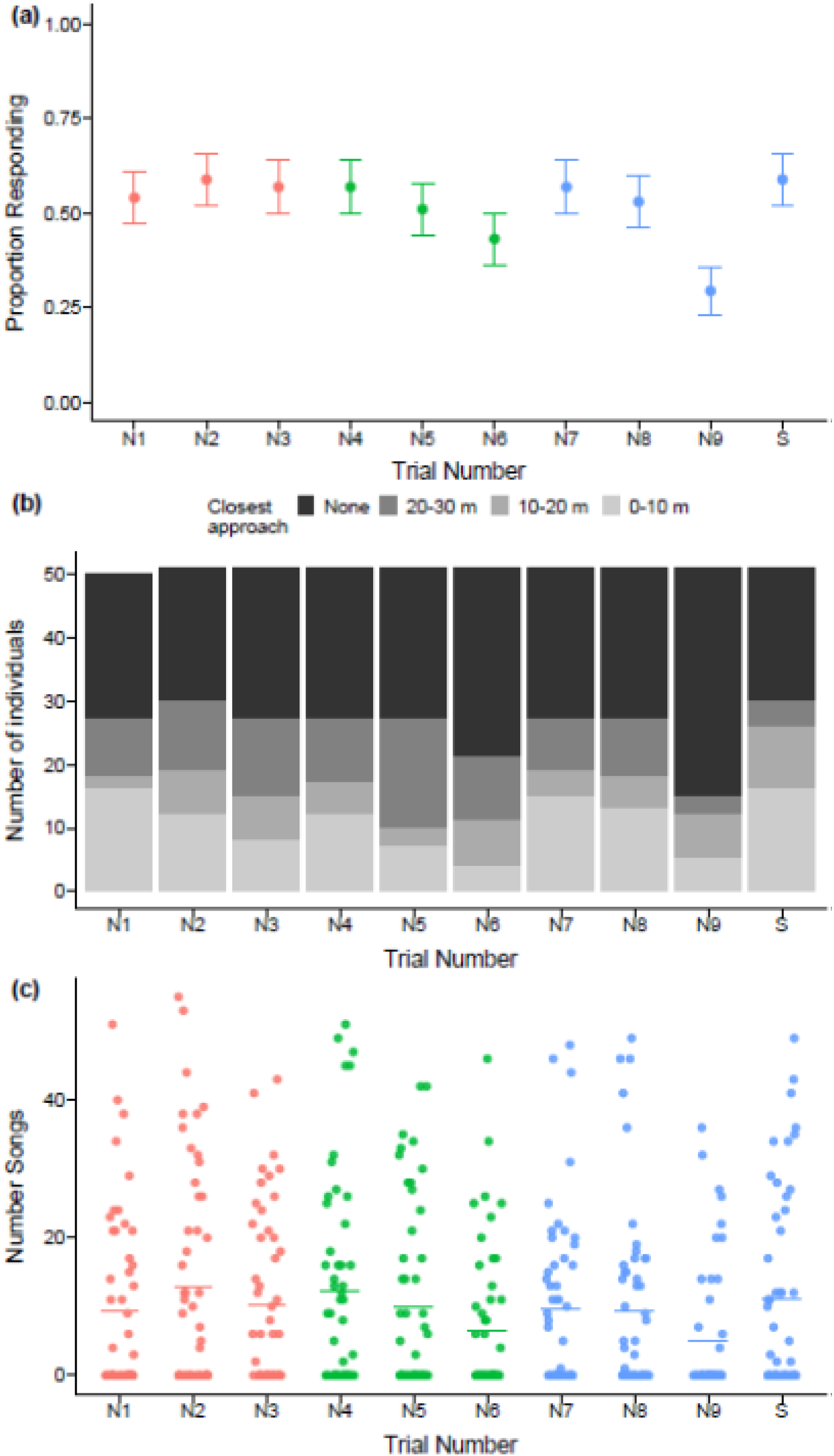
Response across trials. (a) The proportion (±SE) of individuals (*N* = 51 for all trials except *N* = 50 for N1; see Methods) that responded by either singing or approaching the playback speaker in each of the nine neighbour playback trials (labelled N1-N9) and on the stranger playback trial. Colours correspond to the day that the playback was performed (red = day 1, green = day 2, blue = day 3). (b) The closest approach, in categories of 10 m (note that in the statistical analyses we used categories of 5 m, but for ease of visualization we use broader categories here). None corresponds to trials in which the bird did not move at all towards the playback speaker. (c) The number of songs produced by males during the playback trial. Dots represent an individual’s response to that trial (points have been jittered along the x-axis and rendered partially transparent for ease of interpretation). Horizontal lines represent mean values.

### Stimulus specificity of response to neighbour playback

Although twice as many individuals responded to the stranger playback compared to the (immediately preceding) final neighbour playback, this difference was not significant (Fig. 2A; Table 2). However, there was a significant difference in both the closest approach to the speaker and in the number of songs between the final neighbour playback and the stranger playback: individuals approached closer and sang more songs in response to the stranger playback (Fig. 2b, c; Table 2). Thus, the decline in the response was specific to the neighbour’s song (stimulus), which suggests dear enemy discrimination rather than a general decrease in aggression. Stimulus rate did not affect the responses when only these two trials were considered (Table 2).

**Table 2.** The effects of stimulus specificity and stimulus rate on response to neighbour playback. Output is from models as in Table 1, but now only including the response to the final neighbour playback and the stranger playback; estimate shown for the stranger trial (10) and the reference category (estimate = 0) is the final neighbour playback (trial 9). The parameter estimate for the high stimulus rate is shown and the reference category is the low rate. *N* = 51 individuals

| Dependent variable | Factor | Estimate (SE) | z | P |
| --- | --- | --- | --- | --- |
| Response | Intercept | -2.54 (2.51) |  |  |
|  | Stimulus (stranger) | 3.59 (3.33) | 1.08 | 0.28 |
|  | Stimulus rate (high) | -0.12 (1.42) | -0.08 | 0.94 |
| Closest approach | Stimulus (stranger) | -2.09 (0.56) | -3.73 | <0.001 |
|  | Stimulus rate (high) | 0.13 (0.79) | 0.17 | 0.87 |
| Number of songs | Intercept | -0.04 (0.74) |  |  |
|  | Stimulus (stranger) | 0.30 (0.08) | 3.56 | <0.001 |
|  | Stimulus rate (high) | 0.04 (0.92) | 0.04 | 0.97 |

### Repeatability and context dependence of dear enemy behaviour

Based on our standard criterion of no response to the final neighbour playback followed by a response to the stranger playback, 15 of 51 individuals exhibited dear enemy behaviour towards their simulated neighbour after the nine playbacks (Table S2; 15 of 30 when non-responders were removed i.e. standard criterion without non-responders, see Methods; these results refer only to the first set of playbacks, and do not include the second set of playbacks that were performed in 2017). There was a trend for significant repeatability in the standard criterion for dear enemy behaviour, with a moderate repeatability coefficient (*N* = 19 individuals tested twice; R = 0.44, P = 0.074). Five of six individuals that showed dear enemy behaviour in the first set of playbacks also did so in the second, and six of 13 individuals that did not show dear enemy behaviour in the first set of playbacks also did not show it in the second set. There were insufficient numbers of individuals tested twice and responding to the stranger playback to allow for estimating the repeatability of the slope for response or number of songs across trials.

Whether an individual met the standard criterion for dear enemy behaviour (whether including or excluding non-responders) did not depend on the date of testing, the date relative to the day on which the first egg was laid, the year, the average number of songs produced by the subject across the neighbour playbacks, or the stimulus rate (Table S3). These variables also did not affect the slope of the response or number of songs across trials (Table S3). There was also no effect of age (first year versus older) on meeting the standard criterion for dear enemy behaviour (Estimate of effect being older than first year ± SE = −1.29 ± 0.88, z = −1.46, P = 0.15; N = 33 individuals of known age; standard criterion without non-responders: −0.47±0.97, z = −0.48, P = 0.63, N = 18). However, there was an effect of age on the slope of number of songs across trials, with older birds having a less negative slope than first year birds (Estimate = 2.96 ± 0.83, *t* = 3.57, P = 0.003; N = 17 individuals of known age) and also on the slope of responses across trials (Estimate = 0.07 ± 0.03, *t* = 2.33, P = 0.03; N = 17).

### Reproductive success and dear enemy behaviour

There was no effect of whether an individual exhibited dear enemy behaviour (standard criterion) on the number of eggs laid in its nest (Fig. 3a; Table S4). Sixteen offspring were identified as extra-pair young and were excluded from the analyses of number of offspring fledged and offspring mass. There was also no effect of whether an individual exhibited dear enemy behaviour (standard criterion) on the number of within-pair offspring that fledged (Fig. 3b; Table S4), or on the binary variable of whether or not any offspring were fledged (Table S4). For the limited set of individuals that fledged young, there was no effect of whether it had met the standard criterion for dear enemy behaviour on the average mass of its within-pair offspring (Fig. 3c; Table S4) or on the individual mass of within-pair offspring (Table S4). When the standard criterion without non-responders, the slope of responses, or the slope of the number of songs across trials was used as the response variable, there were again no significant effects on any of the reproductive success measures (Table S4). When we repeated the above analyses but this time including data from both within-pair and extra-pair young, we again found no relationship between dear enemy behaviour and any measure of reproductive success (Table S5). Finally, there was no difference between individuals that did or did not show dear enemy behaviour (standard criterion) in the proportion of fledglings that were extrapair (Estimate = 0.20 ± 0.71, P = 0.78, *N* = 29 nests).

**Fig. 3.**
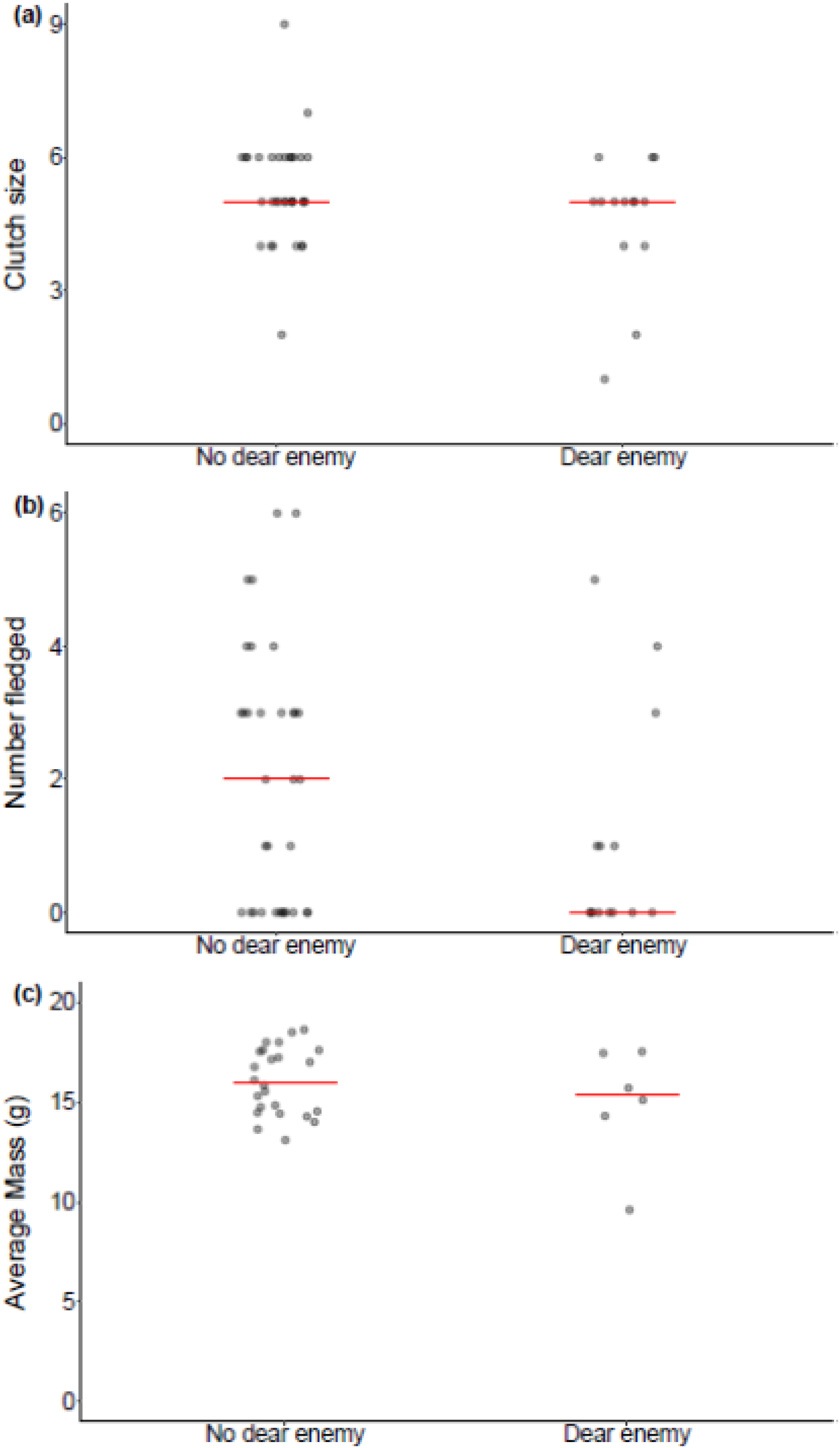
Relationship between whether the individual met the standard criterion (see Methods) for exhibiting the dear enemy effect (“Dear enemy”) or not (“No dear enemy”) and different measures of reproductive success. Each dot represents the value for an individual nest (points jittered along the x-axis and rendered partially transparent); red horizontal line represents the median. (a) Clutch size, (b) Number of within-pair offspring successfully fledged, (c) Average mass of within-pair offspring at day 15 post hatch (not including nests for which no offspring survived to day 15). *N* = 47 individuals in each graph, except for (c) where *N* = 30

## DISCUSSION

Male great tits in our study populations exhibited an overall decline in song response and approach towards the neighbour playback stimulus across repeated presentations, consistent with reduced aggression towards an increasingly familiar individual. This reduced aggression was stimulus-specific because in the stranger playback, simulating a different individual, subjects reverted to a strong response to the playback. This is consistent with dear enemy behaviour because at the end of the trials, individuals were less aggressive towards the neighbour than the stranger. Our experiments show that neighbour-stranger discrimination can arise and be expressed in only three days, although this was not the case for all individuals. Against predictions, there was no fitness benefit for individuals that exhibited dear enemy behaviour, and no reproductive benefit of any kind that we could detect. Below we discuss these results in the context of territorial aggression and the evolution of neighbour-stranger discrimination.

### Learning mechanisms and individual variation

Although complex cognitive mechanisms such as ‘true’ individual recognition (Tibbetts and Dale 2007) are sometimes involved in dear enemy behaviour, and more complex associations between neighbours may arise, perhaps requiring long-term memory, such as coalitions or associations outside the breeding season (McGregor and Avery 1986; Grabowska-Zhang et al. 2012a; Firth and Sheldon 2016), in many species habituation learning of some characteristic of neighbours is a major contributor to the reduced aggression towards those individuals that is characteristic of the dear enemy effect (Petrinovich and Peeke 1973; Brooks and Falls 1975; Peeke 1984; Bee and Gerhardt 2001; Dong and Clayton 2009). We showed that at least some individuals distinguished simulated neighbours from strangers, and that the pattern of response change across trials matches many of the characteristics of habituation (Thompson and Spencer 1966; Rankin et al. 2009).

First, the response to repeated playback of a simulated neighbour decreased across trials (Table 1). Second, there was evidence for spontaneous recovery of the response across days, in which the response on the first trial of the day was generally greater than that of the last trial on the previous day (Fig. 2). A key characteristic of habituation is that over a series of recoveries and decrements, habituation is potentiated, with the response decrement becoming ever more pronounced (Thompson and Spencer 1966; Rankin et al. 2009). Although our nine testing sessions were conducted across just three days, comparisons of behaviour on day 2 and day 3 show that the response decrease was much greater on day 3, consistent with a potentiation of habituation effect. Note that the dear enemy effect does not necessarily require that aggression towards neighbours is eliminated entirely, and indeed increased responsiveness to neighbours at the beginning of the day seems to be common in birds (Briefer et al. 2008; Foote et al. 2008) and may serve to maintain the stability of the territorial boundary. Third, the dual-process theory of habituation argues that in addition to the decrement in response caused by habituation, an independent process of sensitization results in an initial, but transient, increase in aggressive response (Groves and Thompson 1970; Petrinovich and Patterson 1982). Consistent with sensitization, responses tended to be weaker on the first playback trial compared to the subsequent few trials. Importantly, the expression of habituation can be context-dependent through the formation of associations between the habituating stimulus (e.g. the neighbour playback song) and a context (e.g. the location at which the neighbour playback was performed) (Jordan et al. 2000; Uribe-Bahamonde et al. 2019); thus, although our experimental design did not involve independent manipulations of stimulus location and song type, the association between these two variables does not rule out a habituation mechanism for the stimulus-specific reduction in aggression we observed.

Against habituation being the mechanism underlying the observed decline in response, habituation was not stronger at higher song repetition rate, as might have been expected. However, most effects of stimulation interval on habituation have been demonstrated based on variation in the intervals between successive playbacks (rather than between successive songs within a playback), which were approximately the same for the birds in our study, so this prediction may not apply (Groves and Thompson 1970; Thompson et al. 1973; Bee 2001). Taken together, our results suggest that habituation learning is one of the key mechanisms behind the initial stages of dear enemy behaviour in great tits, which was expected, given the general importance of habituation in territorial aggression in songbirds and its specific well-established role in dear enemy recognition (Peeke 1984; Ydenberg et al. 1988; Dong and Clayton 2009). Nevertheless, natural dear enemy relationships in these species almost certainly involve other cognitive mechanisms as well and elucidating the relative contributions of these mechanisms to individual variation in dear enemy behaviour is a worthy target of future research. Furthermore, it is possible that different individuals of the same species use different cognitive mechanisms for individual recognition, and this could have influenced their pattern of response and whether or not they exhibited dear enemy behaviour in this study (Gokcekus et al. 2021).

Non-cognitive mechanisms could also explain the variation in dear enemy behaviour we observed. Differences in average aggressiveness can affect response to playback in great tits (Araya-Ajoy and Dingemanse 2014), and could have played a role, although we found no effect of the average number of songs in response to the neighbour playbacks (a plausible measure of aggressiveness) on whether subjects exhibited dear enemy behaviour. Variation in other personality traits such as explorativeness is also known to relate to variation in habituation speed in great tits (Rivera-Gutierrez et al. 2017) and could therefore have driven some of the variation observed in our study. Given that we did our experiment in a wild population, males could have differed in experience that influenced their response to playbacks. Specifically, males varied in the extent to which they had already interacted with different territorial neighbours in the past. Great tits with more prior exposure to songs of many neighbours are slower to learn to recognize new neighbours (McGregor and Avery 1986), and indeed we found that older birds showed a slower decrement in response and number of songs across the trials than did first year birds, although this did not result in differences between the age groups in whether they differed in response to the final neighbour playback versus the stranger playback. Factors such as density and progress of the breeding season often explain variation in the dear enemy effect (Hyman 2005; Pratt and McLain 2006; Yoon et al. 2012), including in great tits (Jin et al. 2021). However, we found no evidence for this in our study: there was no effect of year, date, or date relative to the start of hatching on whether individuals expressed the dear enemy effect. Whatever the exact mechanisms involved, the individual repeatability in dear enemy behaviour we report indicates that we captured intrinsic differences among individuals in how they discriminated between familiar neighbours and strangers.

We found behaviour consistent with the dear enemy effect because over time individuals responded less to the stimulus simulating the establishment of a new neighbour than to a stimulus that simulated an unknown stranger. It could be objected that subjects did not actually recognize the playback as a true neighbour, and the other as a different stranger bird, and that therefore the stimulus-specific response decrement we observed is not the same as the dear enemy effect. In the strict sense, even playback designs using songs of true neighbours do not necessarily overcome this objection, because even demonstrating a stimulus- and location-specific response to a true neighbour’s songs does not give definitive evidence that this stimulus was recognized as the territorial neighbour, rather than being a stimulus that the individual had simply habituated to. We justify our approach and interpretation because 1) we were interested in individual variation in the cognitive processes involved in establishing dear enemy relationships, which cannot be studied with established neighbours. Given the frequency of habituation as a mechanism for dear enemy recognition and the very rapid settlement of territorial relations and habituation to song in great tits (Krebs et al. 1981; Peeke 1984; Rivera-Gutierrez et al. 2017), our design was appropriate. 2) Although no playback can ever replicate the interactions that take place between two birds, we replicated the essential features of territorial neighbours – an increasingly familiar song from the same location on an adjacent territory – and those of strangers – an unfamiliar song from a different location. Song familiarity and location are likely the two most important elements in dear enemy behaviour in great tits, and the behaviour we observed was consistent with the expression of the dear enemy effect. 3) Our simulated neighbour approach reduces some of the confounding effects associated with using playbacks of an established neighbour such as individual variation in experience with that neighbour (Falls et al. 1982; Grabowska-Zhang et al. 2012a). Nevertheless, additional studies of dear enemy behaviour in this species that examine its expression over longer time periods and incorporating additional stimulus dimensions and long-term observations of interacting individuals would be valuable.

### Dear enemy recognition and fitness

We hypothesized that if the dear enemy effect is adaptive because it reduces time spent in unnecessary aggressive interactions or leads to other beneficial interactions with neighbours (Getty 1987; Temeles 1994), then individuals that reduce the aggression directed towards their neighbours more quickly would have higher reproductive benefits leading to increased reproductive fitness. There was good reason to expect these predictions to be met in our system because male great tits are known to trade off foraging and territory defence (Ydenberg 1984). We reiterate that our experiment was based on the premise that individual variation in the reduction in aggression to the neighbour stimulus and subsequent response to the stranger stimulus served as a proxy for individual variation in the tendency to form dear enemy relationships in natural interactions, and it was the latter that we proposed to drive any fitness effects. Our hypothesis was not supported because there was no difference in clutch size, average biomass, or number of offspring that fledged between individuals that did or did not express the dear enemy effect. This was true in terms of the measure most closely related to male fitness (when extra-pair paternity was excluded) but it was also true for the overall reproductive success at the nest (when extra-pair young were included), for which detectable effects should be most apparent. In a field experiment of this nature on a species with a complex multi-stage reproductive strategy, inevitably many processes could explain this result, and we highlight those that we suggest particularly deserve attention in future research.

First, ‘enemies may not always be dear’ (Muller and Manser 2007; Courvoisier et al. 2014; Moser-Purdy et al. 2017), and there may be costs associated with reducing aggression. For example it may lead to prospective individuals settling, leading to reduced territory size and quality (Getty 1989), or to reduced rates of provisioning offspring by the males (Sillett et al. 2004). Reduced aggression by males could have also led to the cost of their mates engaging in extrapair copulations with neighbours, though we found no evidence of this, nor did a previous study of song sparrows, *Melospiza melodia* (Krippel et al. 2017). We note, however, that reduced aggression could also have led to reduced extra-pair paternity if it facilitated mate guarding, though we also found no evidence for this. Second, the lack of any observed benefit to dear enemy behaviour could be because it traded off against other fitness-related behaviours. For instance, in great tits, a positive association between problem solving performance and both clutch size and number of fledglings was masked by a positive association with nest desertion leading to no net reproductive benefits (Cole et al. 2012). We observed high levels of nest failure due to either abandonment or natural death of offspring among birds that expressed dear enemy behaviour, which may be an indication of such a cost, although the likelihood of nest failure was not significantly different from birds that did not express dear enemy behaviour. However, perhaps a more likely reason for no observed effect is that our fitness measures were for a single breeding season, but typically selection varies over time (Siepielski et al. 2009). Similarly, we examined the initial stages of formation of dear enemy relationships and it may be that variation in the strength and specificity of relationships with neighbours over the course of the whole breeding season is more consequential for fitness, although these temporal dynamics and their relationship with fitness have not been explored. Third, although our measures of dear enemy behaviour had a high (albeit non-significant) repeatability coefficient, suggesting the possibility of intrinsic differences among individuals, in reality our measure was likely to have been influenced by other factors—for example temporary environmental effects—making it more difficult to identify links between dear enemy behaviour and variation in fitness (Zsebők et al. 2017). And finally, alterations to the experimental design, such as using a taxidermic mount to simulate visual cues (Ritschard et al. 2012; Araya-Ajoy et al. 2016) or carrying out the playbacks for more than three days, may have revealed additional individual variation that is more directly linked to dear enemy behaviour.

Dear enemy behaviours are undoubtedly adaptive and may well be under current selection in many systems, especially in species such as the great tit where territorial behaviour is important (Krebs 1971; Falls et al. 1982; McGregor and Avery 1986; Grabowska-Zhang et al. 2012b). Despite the intrinsic differences among individuals in their tendency to reduce the response to a familiar stimulus and then respond to a novel stimulus, none of these behaviours showed any positive or negative association with a variety of life history traits. However, revealing hypothetical links on behavioural traits such as the dear enemy effect that are likely underpinned by a variety of mechanisms, including cognitive mechanisms that themselves influence other functional behaviours, is likely to be more challenging than for behaviours more closely related to reproduction, and to require a multivariate and longer-term approach across different ecological conditions (Morand-Ferron et al. 2016).

## Acknowledgments

Karen Cogan assisted with equipment sourcing and purchasing. Maike Foraita assisted with playback trials. Will O’Shea, Paul Whitelaw, Sam Bayley, Iván de la Hera, Amy Cooke, and Jenny Coomes helped with nestbox monitoring during the breeding season. Three anonymous reviewers provided helpful comments on a previous draft of this article.

## Author contributions

MSR and JLQ conceived of the project, MSR, JMSC and JON performed the playbacks, MSR, GLD, IGK, JMC and JON collected breeding data, ED, CS and KvO did the DNA extractions and analyses of paternity, MSR analyzed the data, and MSR wrote the manuscript with input from all authors. Note that middle authors on this manuscript are listed in alphabetical order by surname.

## Data availability statement

Raw data associated with this project are available at https://doi.org/10.6084/m9.figshare.17148338.v1.

## Ethical Approval

The project received ethical approval from the Animal Welfare Body at University College Cork (HPRA license number AE19130-P017) and was in accordance with the ASAB (Association for the Study of Animal Behaviour) Guidelines for the Treatment of Animals in Behavioural Research and Teaching. All research was conducted under licenses from the National Parks and Wildlife Service of Ireland and the British Trust for Ornithology as part of ongoing research in these populations.

---

**Table S1.**
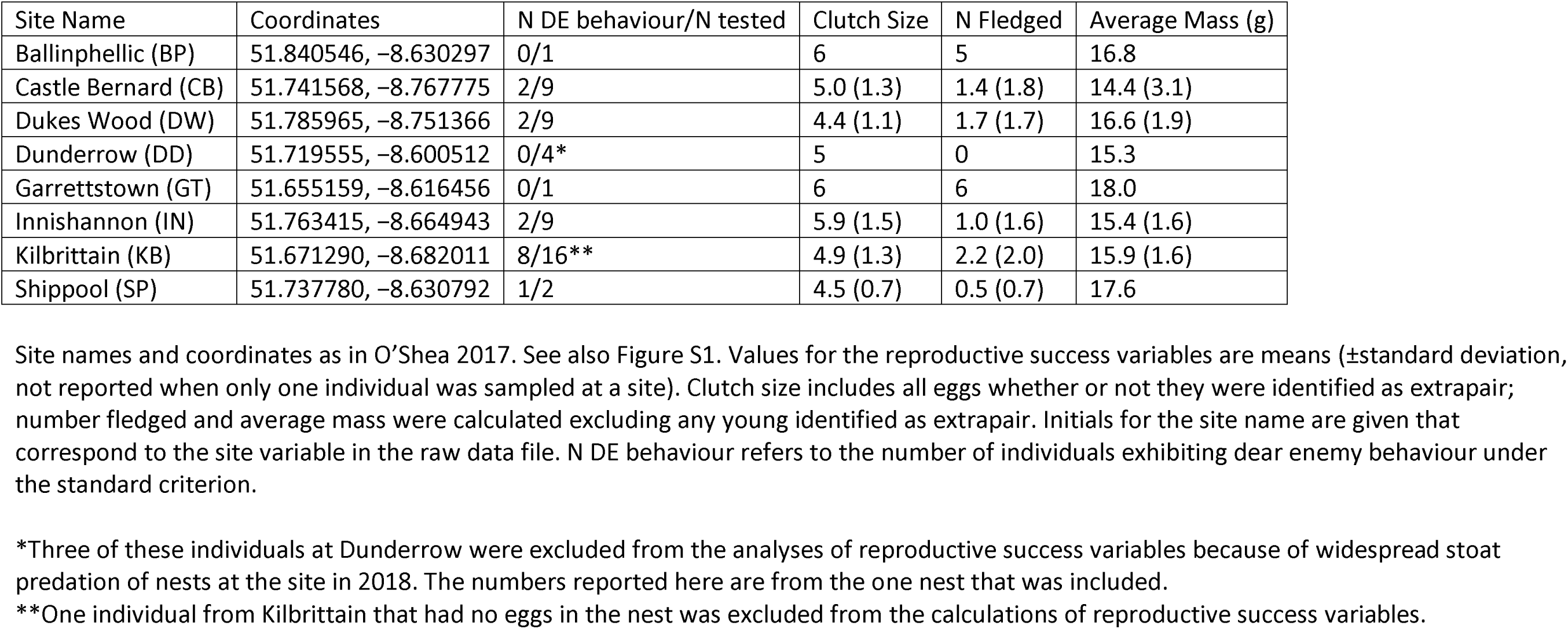
Study site locations and summary statistics.

| Site Name | Coordinates | N DE behaviour/N tested | Clutch Size | N Fledged | Average Mass (g) |
| --- | --- | --- | --- | --- | --- |
| Ballinphellic (BP) | 51.840546, -8.630297 | 0/1 | 6 | 5 | 16.8 |
| Castle Bernard (CB) | 51.741568, -8.767775 | 2/9 | 5.0 (1.3) | 1.4 (1.8) | 14.4 (3.1) |
| Dukes Wood (DW) | 51.785965, -8.751366 | 2/9 | 4.4 (1.1) | 1.7 (1.7) | 16.6 (1.9) |
| Dunderrow (DD) | 51.719555, -8.600512 | 0/4* | 5 | 0 | 15.3 |
| Garrettstown (GT) | 51.655159, -8.616456 | 0/1 | 6 | 6 | 18.0 |
| Innishannon (IN) | 51.763415, -8.664943 | 2/9 | 5.9 (1.5) | 1.0 (1.6) | 15.4 (1.6) |
| Kilbritten (KB) | 51.671290, -8.682011 | 8/16** | 4.9 (1.3) | 2.2 (2.0) | 15.9 (1.6) |
| Shippool (SP) | 51.737780, -8.630792 | 1/2 | 4.5 (0.7) | 0.5 (0.7) | 17.6 |

\*Three of these individuals at Dunderrow were excluded from the analyses of reproductive success variables because of widespread stoat predation of nests at the site in 2018. The numbers reported here are from the one nest that was included.
\*\*One individual from Kilbritten that had no eggs in the nest was excluded from the calculations of reproductive success variables.

**Figure S1.**
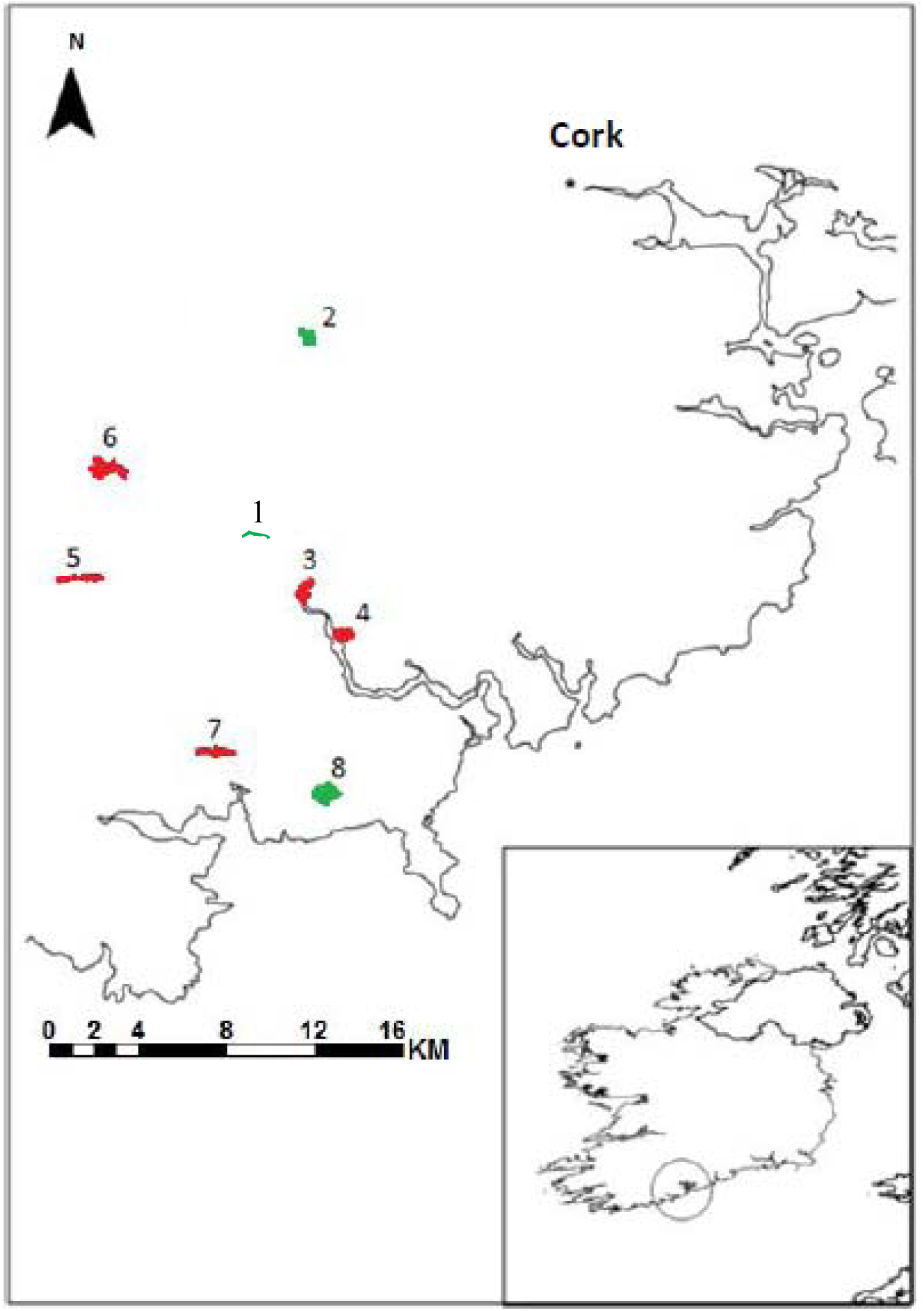
Map of the study sites, modified from O’Shea (2017). Numbers correspond to the individual sites (see also Table S1): 1. Innishannon, 2. Ballinphellic, 3. Shippool, 4. Dunderrow, 5. Castle Bernard, 6. Dukes Wood, 7. Kilbrittain, 8. Garrettstown.

**Table S2.**
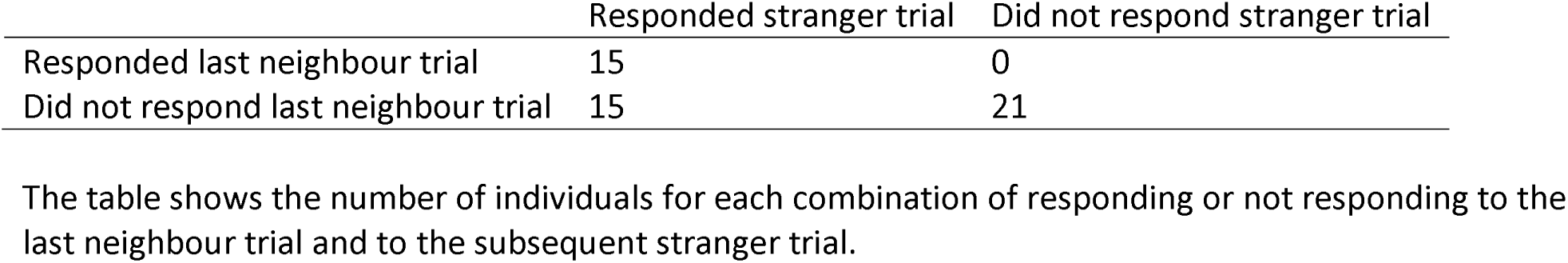
Response on last neighbour trial versus response on stranger trial

**Table S3.** Effects of context on dear enemy behaviour.

| Dear enemy criterion | Factor | Estimate | SE | Test statistic | P |
| --- | --- | --- | --- | --- | --- |
| Standard criterion | Intercept | 1049 | 1668 | 0.63 | 0.53 |
|  | Year | -0.52 | 0.83 | -0.63 | 0.53 |
|  | Date | 0.07 | 0.05 | 1.33 | 0.18 |
|  | Stimulus rate (high) | 0.58 | 0.70 | 0.82 | 0.41 |
|  | Date relative to first egg | -0.01 | 0.06 | -0.18 | 0.86 |
|  | Average song number | -0.07 | 0.06 | -1.13 | 0.26 |
| Standard without nonresponders | Intercept | 5701 | 3478 | 1.64 | 0.10 |
|  | Year | -2.83 | 1.73 | -1.64 | 0.10 |
|  | Date | 0.15 | 0.10 | 1.52 | 0.13 |
|  | Stimulus rate (high) | 0.87 | 1.08 | 0.81 | 0.42 |
|  | Date relative to first egg | -0.02 | 0.08 | -0.26 | 0.80 |
|  | Average song number | -0.25 | 0.12 | -2.09 | 0.04 |
| Response slope | Intercept | 12.37 | 73.49 | 0.17 | 0.87 |
|  | Year | -0.01 | 0.04 | -0.17 | 0.87 |
|  | Date | 0.001 | 0.002 | 0.46 | 0.65 |
|  | Stimulus rate (high) | 0.01 | 0.03 | 0.29 | 0.77 |
|  | Date relative to first egg | 0.002 | 0.002 | 1.05 | 0.3 |
|  | Average song number | 0.002 | 0.002 | 1.00 | 0.33 |
| Song slope | Intercept | 485 | 2262 | 0.21 | 0.83 |
|  | Year | -0.24 | 1.12 | -0.22 | 0.83 |
|  | Date | 0.08 | 0.07 | 1.13 | 0.27 |
|  | Stimulus rate (high) | -0.70 | 0.81 | -0.86 | 0.40 |
|  | Date relative to first egg | 0.01 | 0.06 | 0.12 | 0.91 |
|  | Average song number | 0.05 | 0.05 | 0.96 | 0.35 |

Results are from a generalized linear model of whether or not the individual met the standard criterion for expressing dear enemy behaviour (where the test statistic is $z$ ), or from linear models for the slope variables (where the test statistic is $t$ ). The reference category for stimulus rate is the low rate stimulus. The variable “average song number” refers to the average number of songs produced by the bird over the nine total neighbour trials. Date is coded as the day number of the year on which the first playback was performed for that bird, with 1 March being day 1. Date relative to first egg refers to which day the first playback was performed for that bird relative to when the first egg was laid in the nest (so day zero would indicate that the first playback was performed on the same day that the first egg was laid in the nest). $N = 50$ individuals for the standard criterion (one individual not included because there was never an egg laid at the nest), 29 individuals for standard without nonresponders, and 25 individuals for the slope variables.

**Table S4.**
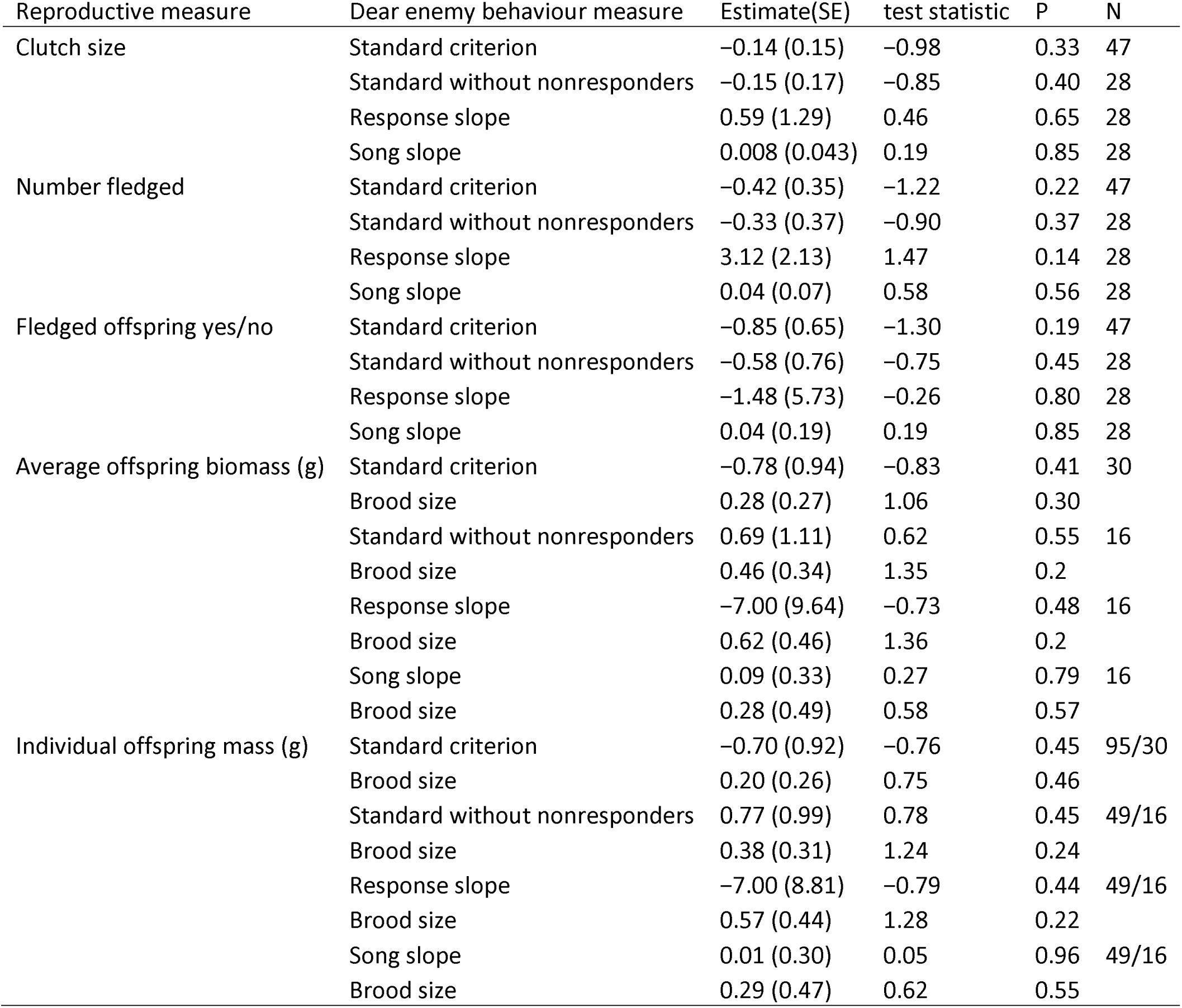
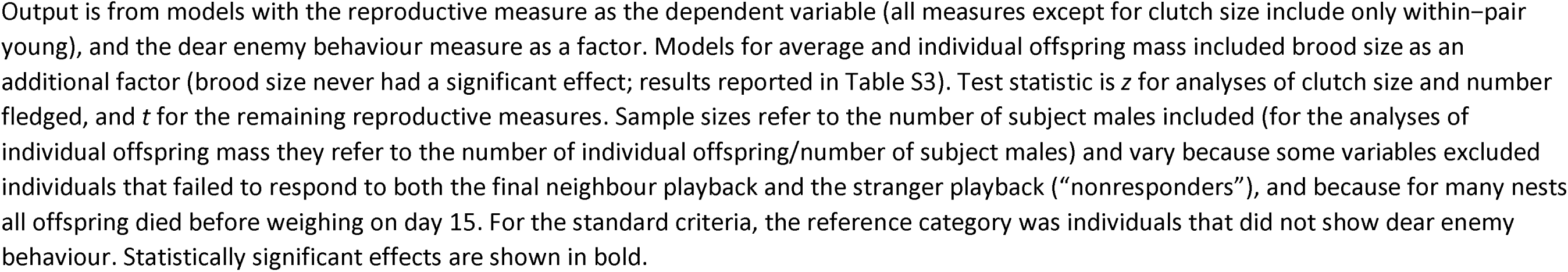
Full models of dear enemy behaviour and reproductive success, including brood size effects when included in models.

| Reproductive measure | Dear enemy behaviour measure | Estimate(SE) | test statistic | P | N |
| --- | --- | --- | --- | --- | --- |
| Clutch size | Standard criterion | -0.14 (0.15) | -0.98 | 0.33 | 47 |
|  | Standard without nonresponders | -0.15 (0.17) | -0.85 | 0.40 | 28 |
|  | Response slope | 0.59 (1.29) | 0.46 | 0.65 | 28 |
|  | Song slope | 0.008 (0.043) | 0.19 | 0.85 | 28 |
| Number fledged | Standard criterion | -0.42 (0.35) | -1.22 | 0.22 | 47 |
|  | Standard without nonresponders | -0.33 (0.37) | -0.90 | 0.37 | 28 |
|  | Response slope | 3.12 (2.13) | 1.47 | 0.14 | 28 |
|  | Song slope | 0.04 (0.07) | 0.58 | 0.56 | 28 |
| Fledged offspring yes/no | Standard criterion | -0.85 (0.65) | -1.30 | 0.19 | 47 |
|  | Standard without nonresponders | -0.58 (0.76) | -0.75 | 0.45 | 28 |
|  | Response slope | -1.48 (5.73) | -0.26 | 0.80 | 28 |
|  | Song slope | 0.04 (0.19) | 0.19 | 0.85 | 28 |
| Average offspring biomass (g) | Standard criterion | -0.78 (0.94) | -0.83 | 0.41 | 30 |
|  | Brood size | 0.28 (0.27) | 1.06 | 0.30 |  |
|  | Standard without nonresponders | 0.69 (1.11) | 0.62 | 0.55 | 16 |
|  | Brood size | 0.46 (0.34) | 1.35 | 0.2 |  |
|  | Response slope | -7.00 (9.64) | -0.73 | 0.48 | 16 |
|  | Brood size | 0.62 (0.46) | 1.36 | 0.2 |  |
|  | Song slope | 0.09 (0.33) | 0.27 | 0.79 | 16 |
|  | Brood size | 0.28 (0.49) | 0.58 | 0.57 |  |
| Individual offspring mass (g) | Standard criterion | -0.70 (0.92) | -0.76 | 0.45 | 95/30 |
|  | Brood size | 0.20 (0.26) | 0.75 | 0.46 |  |
|  | Standard without nonresponders | 0.77 (0.99) | 0.78 | 0.45 | 49/16 |
|  | Brood size | 0.38 (0.31) | 1.24 | 0.24 |  |
|  | Response slope | -7.00 (8.81) | -0.79 | 0.44 | 49/16 |
|  | Brood size | 0.57 (0.44) | 1.28 | 0.22 |  |
|  | Song slope | 0.01 (0.30) | 0.05 | 0.96 | 49/16 |
|  | Brood size | 0.29 (0.47) | 0.62 | 0.55 |  |

**Table S5.**
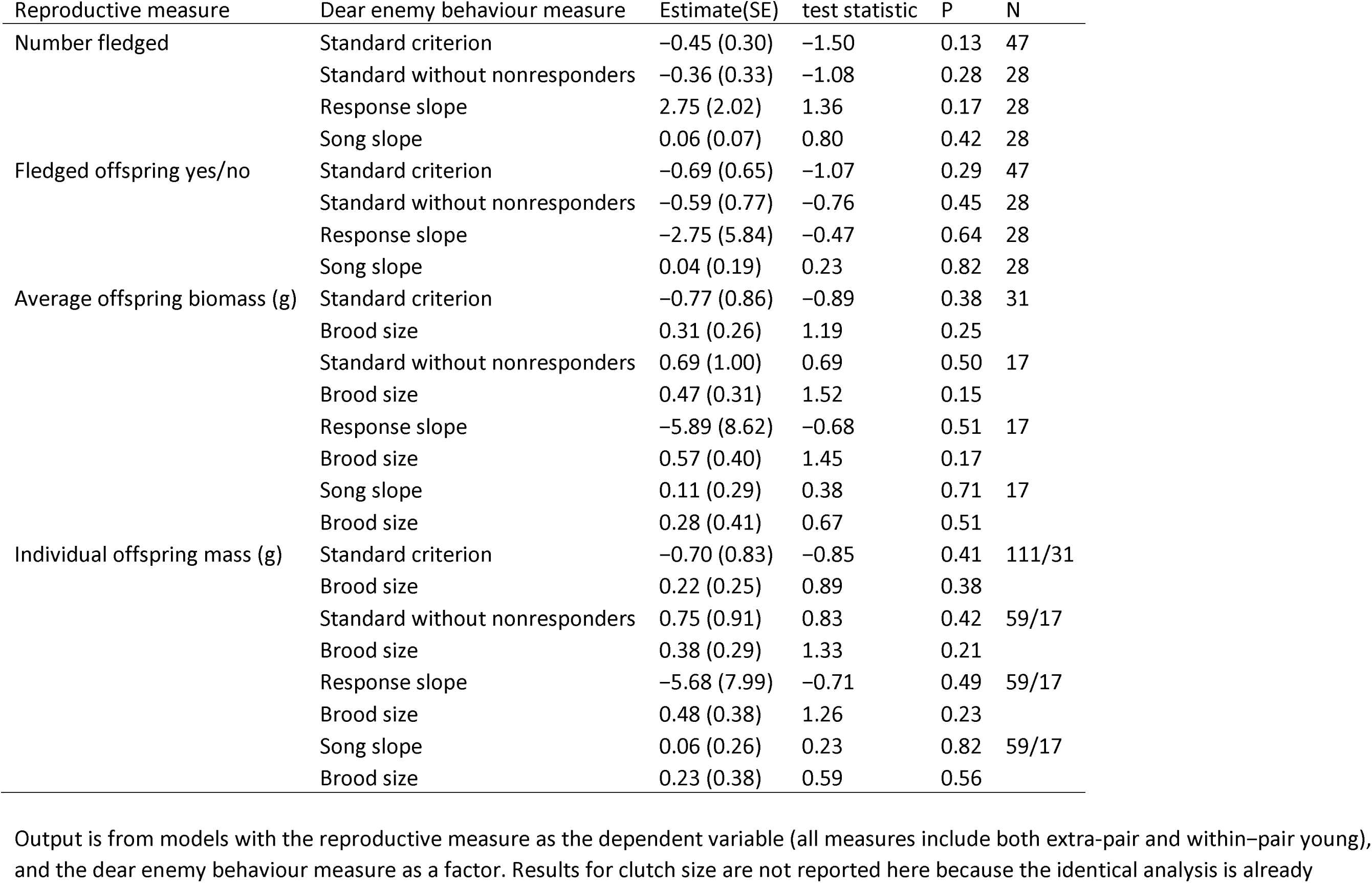
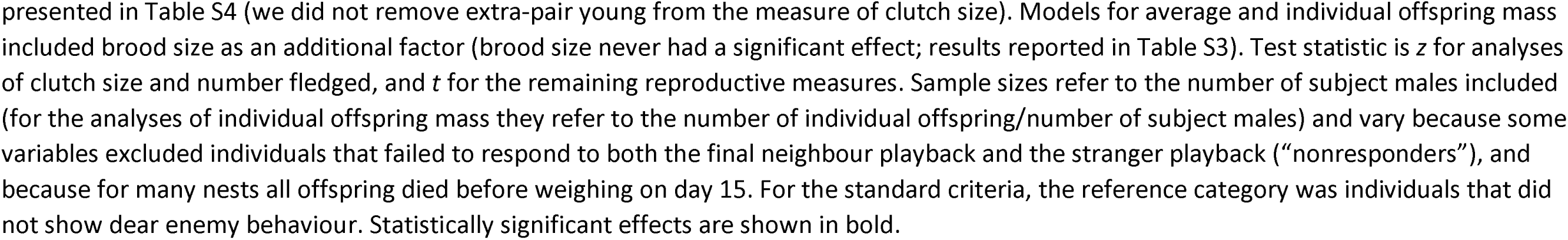
Full models of dear enemy behaviour and reproductive success. Unlike Table S4, these analyses include both within-pair and extra-pair young.

